## Supplemental figures for "circZNF827 nucleates a transcription inhibitory complex to balance neuronal differentiation"

Figure 1 – figure supplement 1

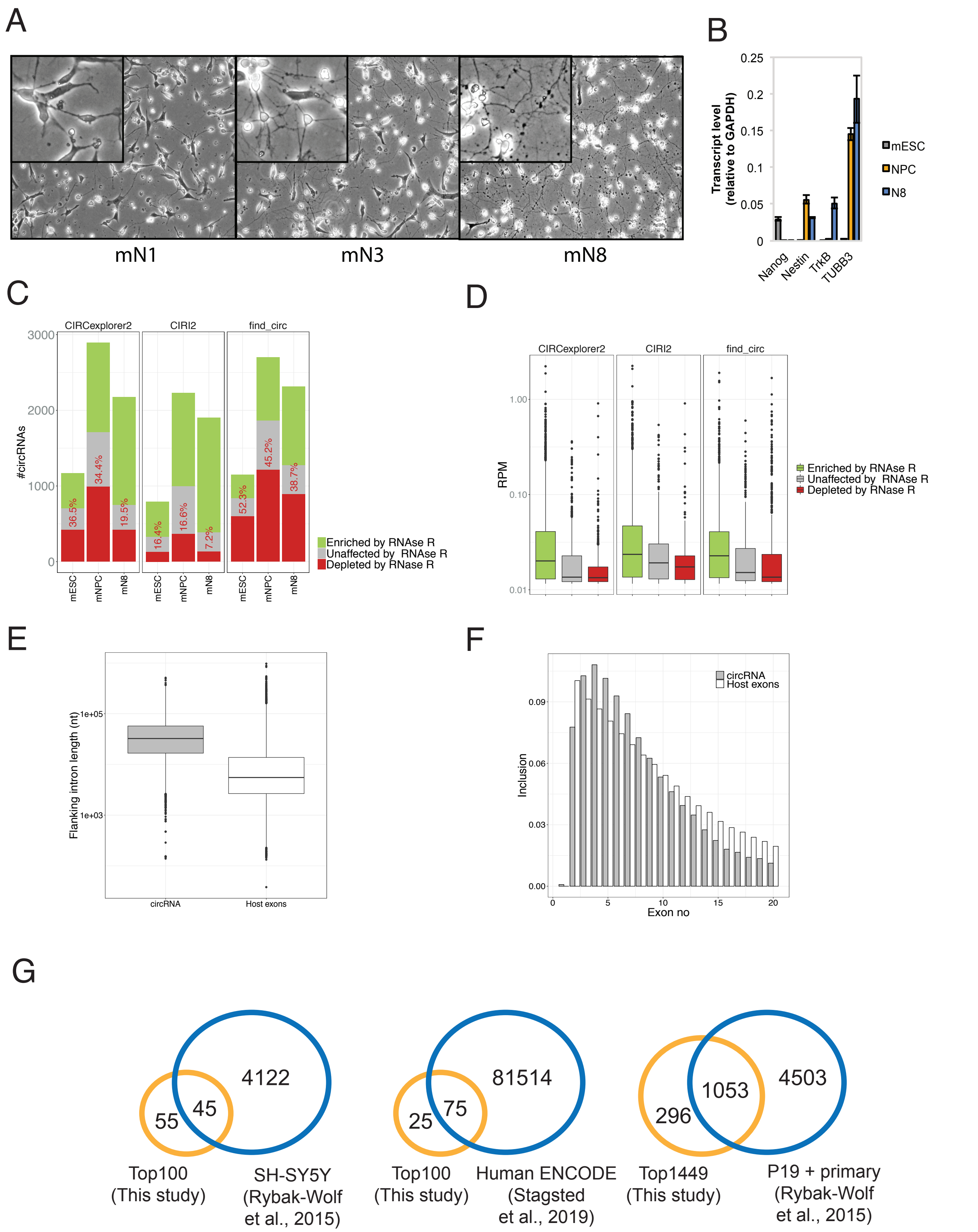

Figure 2 – figure supplement 1

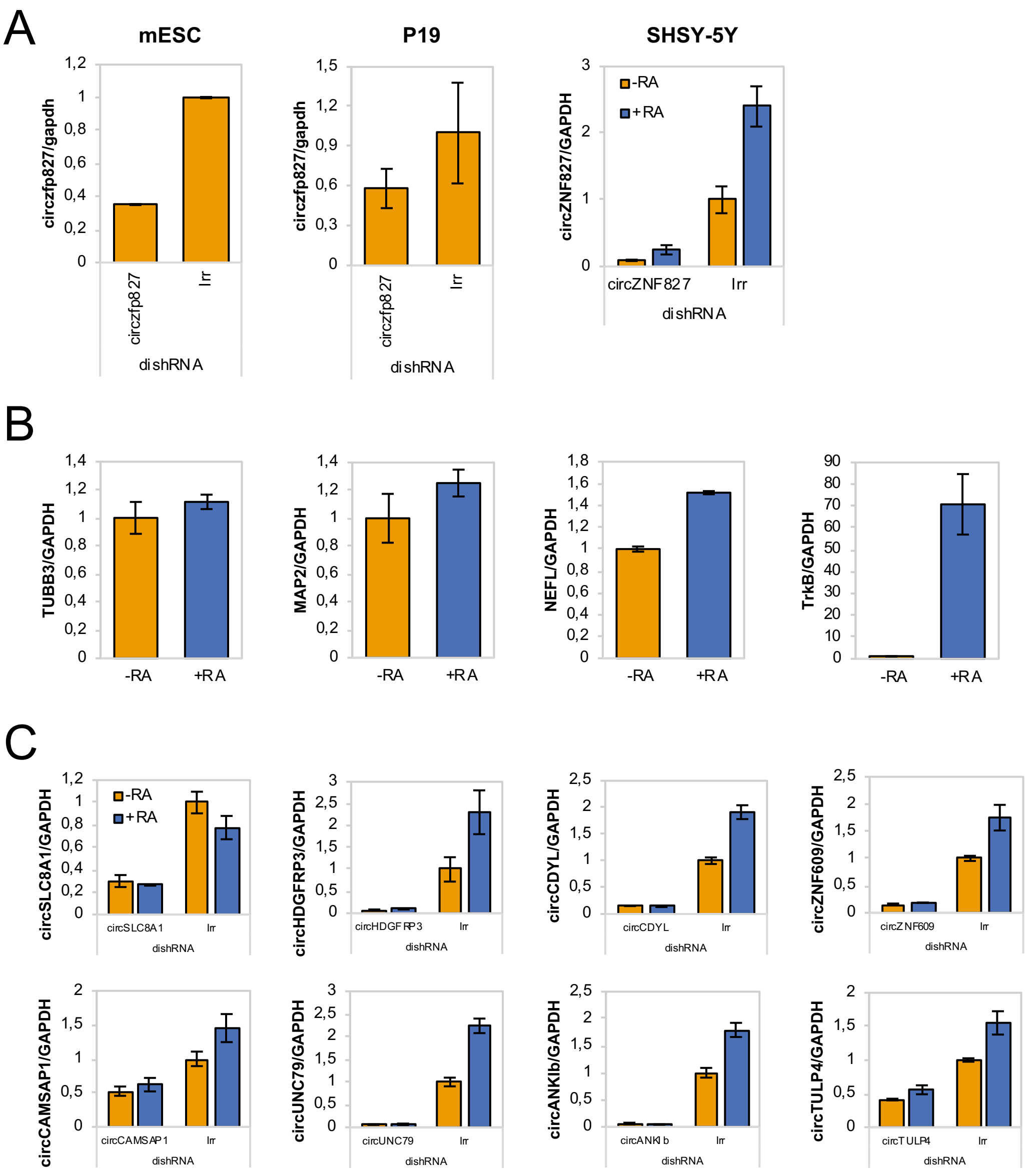


Hollensen et al., 2020

Figure 2 – figure supplement 2

A

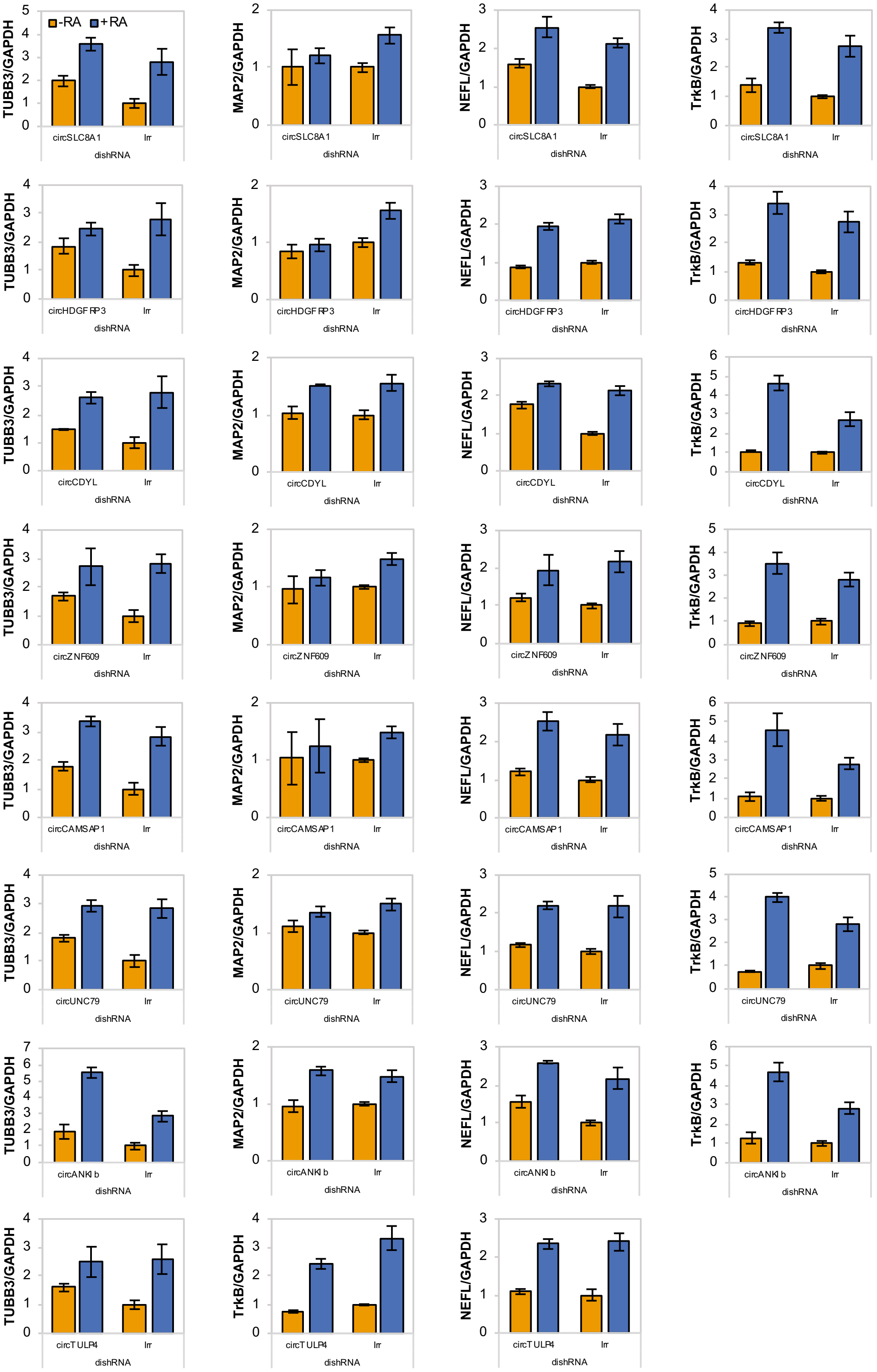


B

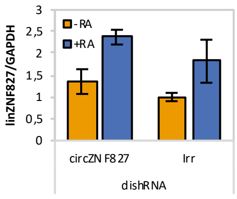


Hollensen et al., 2020

### Figure 2 - figure supplement 3

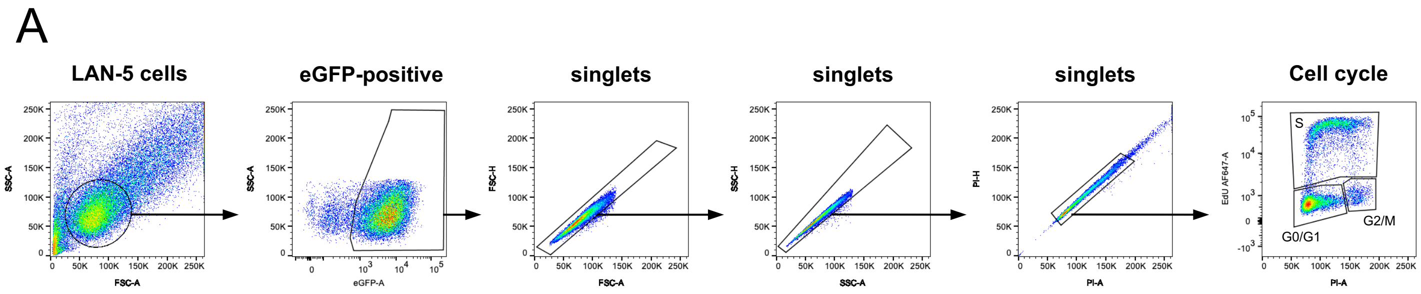


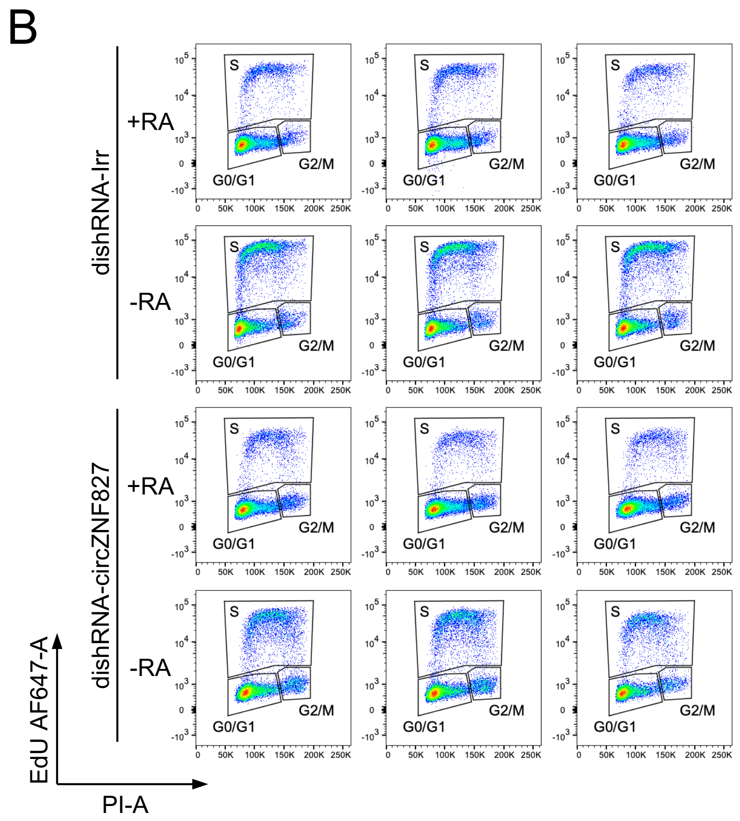


Hollensen et al., 2020

Figure 3 - figure supplement 1

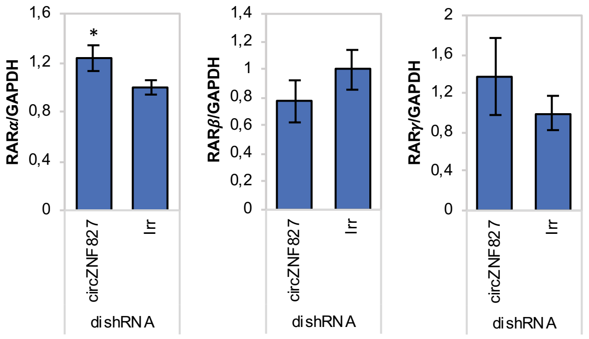


Hollensen et al., 2020

Figure 4 - figure supplement 1

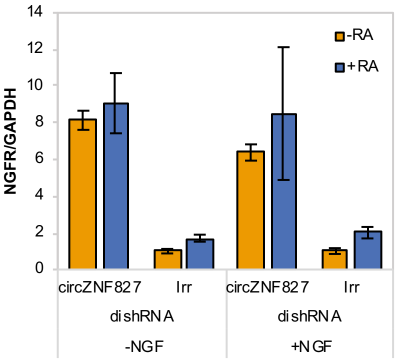


Hollensen et al., 2020

### Figure 5 – figure supplement 1

A

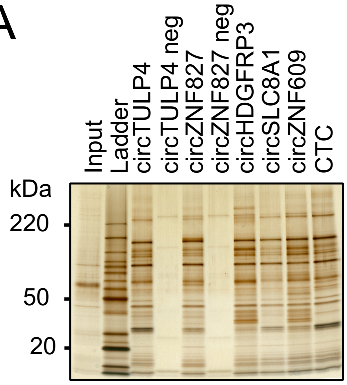


C

hsa_circZNF827:
BSJ_CGGAAAGCTACAAGGAAACACAGATGGTGAAGATTAAAGAGGAACCCATGGAGGTT
GACATCCAGGACTCCCATGTCTCGATATCACCCAGCCGGAATGTTGGCTACAGCACTTTA
ATCGGGCGAGAGAAAACCGAACCCTTACAGAAGATGCCAGAGGGCAGAGTACCCCCAGAG
AGAAACCTCTTCAGTCAGGATATCTCTGTGAAAATGGCTTCCGAGCTCCTCTTTCAACTG
TCAGAAAAAGTGAGCAAAGAGCACAATCATACAAAAGAAAACACCATCCGGACCACGACC
AGCCCTTTCTTTTCAGAAGACACATTTAGACAATCACCATTCACCTCCAATTCAAAAGAA
CTGCTGCCCAGTGACTCCGTGCTGCACGGAAGAATATCAGCTCCAG_BSJ

Putative binding sites: hnRNP K and hnRNP L

B

| Protein: HNRNPK(Hs/Mm) | | | | |
|---|---|---|---|---|
| Position | Motif | Occurrence | Z-score | P-value |
| 295 | ccawmcc | aaagaaaacaccauccggaccacgaccagcccuuucuuuucagaagacacauuuaga | 1.739 | 4.10e-02 |
| Protein: HNRNPL(Hs/Mm) | | | | |
| Position | Motif | Occurrence | Z-score | P-value |
| 257 | acacrav | aacugucagaaaaagugagcaaagagcacaaucauacaaaagaaaacaccauccgga | 2.887 | 1.94e-03 |
| 259 | amayama | cugucagaaaaagugagcaaagagcacaaucauacaaaagaaaacaccauccggacc | 2.813 | 2.45e-03 |
| 263 | amayama | cagaaaaagugagcaaagagcacaaucauacaaagaaaacaccauccggaccacga | 2.987 | 1.41e-03 |
| 265 | acacrav | gaaaaagugagcaaagagcacaaucauacaaaagaaaacaccauccggaccacgacc | 3.394 | 3.44e-04 |
| 267 | acacrav | aaaagugagcaaagagcacaaucauacaaaagaaaacaccauccggaccacgaccag | 3.465 | 2.65e-04 |
| 275 | amayama | gcaaagagcacaaucauacaaaagaaaacaccauccggaccacgaccagcccuuucu | 3.093 | 9.91e-04 |
| 277 | acacrav | aaagagcacaaucauacaaaagaaaacaccauccggaccacgaccagcccuuucuuu | 3.563 | 1.83e-04 |
| 289 | acacrav | cauacaaaagaaaacaccauccggaccacgaccagcccuuucuuuucagaagacaca | 2.972 | 1.48e-03 |
| 314 | amayama | ccacgaccagcccuuucuuuucagaagacacauuuagacaaucaccauucaccucca | 2.480 | 6.57e-03 |
| 316 | amayama | acgaccagcccuuucuuuucagaagacacauuuagacaaucaccauucaccuccaau | 2.133 | 1.65e-02 |
| 326 | amayama | cuuucuuuucagaagacacauuuagacaaucaccauucaccuccaauucaaaagaac | 2.160 | 1.54e-02 |

RBPmap precitions, http://rbpmap.technion.ac.il/

D

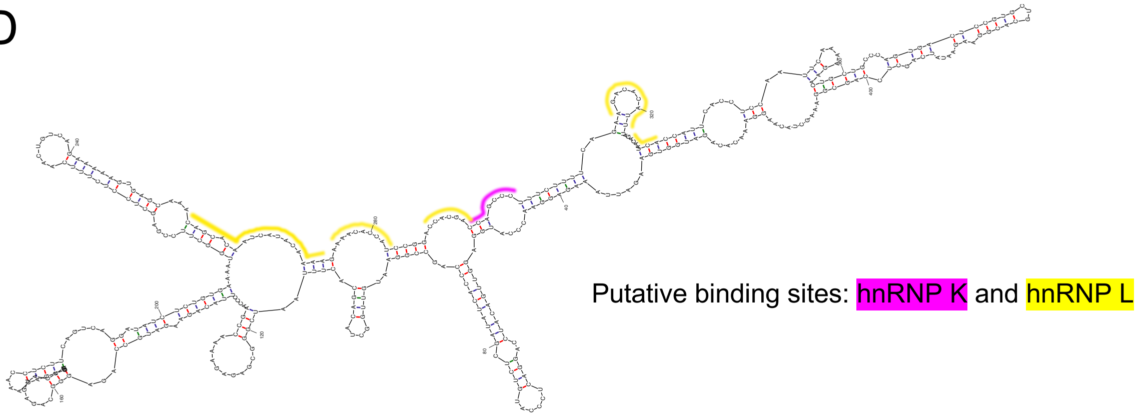


E

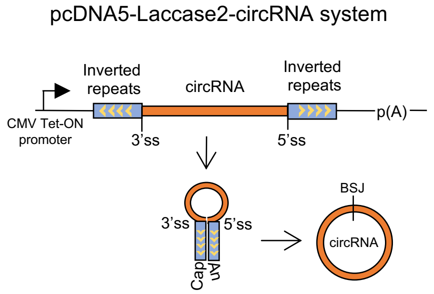


F

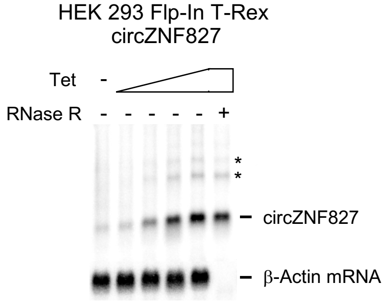


Hollensen et al., 2020

### Figure 6 – figure supplement 1

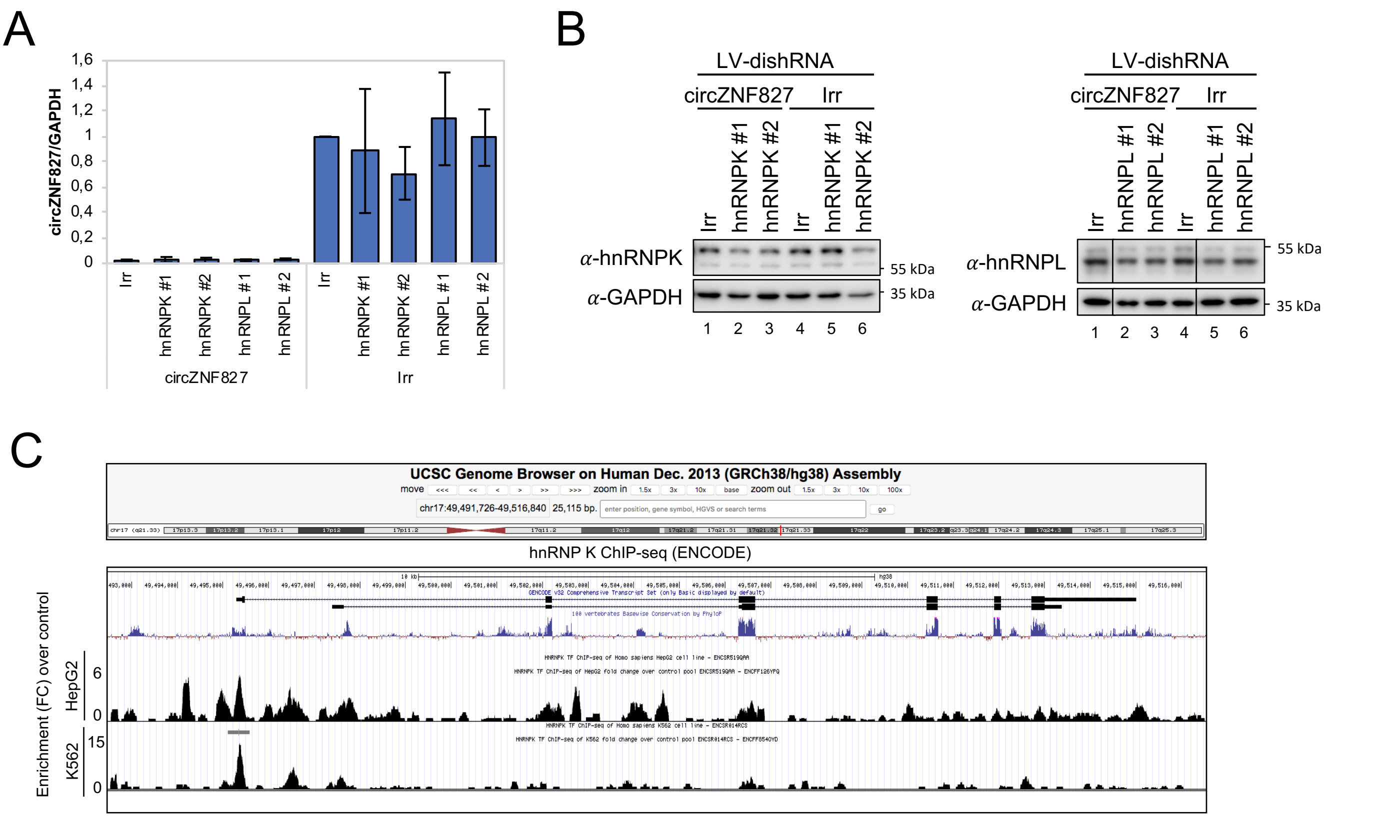
