## Supplemental tables for "circZNF827 nucleates a transcription inhibitory complex to balance neuronal differentiation"

Table S1

See Supplementary Table S1 (excel file)

### Table S2

| Human | | | | | | Mouse | | | | | |
|---|---|---|---|---|---|---|---|---|---|---|---|
| **circRNA** | GRCh37/hg19 | | | circBase ID | Gene | **circRNA** | GRCm38/mm10 | | | circBase ID | Gene |
| | Chromosome | Start | End | | | | Chromosome | Start | End | | |
| **circANKIb** | chr7 | 91924202 | 91957214 | hsa_circ_0135062 | ANKIB1 | **circAnkib1** | chr5 | 3747020 | 3772787 | mmu_circ_0001311 | Ankib1 |
| **circCAMSAP1** | chr9 | 138773478 | 138774924 | hsa_circ_0001900 | CAMSAP1 | **circCamsap1** | chr2 | 25821182 | 25822468 | mmu_circ_0009762 | Camsap1 |
| **circCDYL** | chr6 | 4891946 | 4892613 | hsa_circ_0008285 | CDYL | **circCdyl** | chr13 | 35907777 | 35908444 | mmu_circ_0000451 | Cdyl |
| **circHDGFRP3** | chr15 | 83819966 | 83832827 | hsa_circ_0000646 | HDGFRP3 | **circHdgfrp3** | chr7 | 89038684 | 89050522 | mmu_circ_0001589 | Hdgfrp3 |
| **circHIPK3** | chr11 | 33307958 | 33309057 | hsa_circ_0000284 | HIPK3 | **circHipk3** | chr2 | 104310905 | 104312004 | mmu_circ_0001052 | Hipk3 |
| **circMAGI** | chr3 | 65464266 | 65479306 | hsa_circ_0066459 | MAGI1 | **circMagi1** | chr6 | 93742340 | 93765819 | mmu_circ_0001496 | Magi1 |
| **circMED13L** | chr12 | 116675272 | 116675510 | hsa_circ_0028587 | MED13L | **circMed13l** | chr5 | 119043341 | 119043579 | mmu_circ_0001396 | Med13l |
| **circNFIX** | chr19 | 13135834 | 13136366 | hsa_circ_0005660 | NFIX | **circNfix** | chr8 | 87295682 | 87296214 | mmu_circ_0001704 | Nfix |
| **circRMST** | chr12 | 97886238 | 97954825 | hsa_circ_0099634 | RMST | **circRmst** | chr10 | 91537914 | 91599435 | mmu_circ_0000205 | Rmst |
| **circSLC8A1** | chr2 | 40655612 | 40657441 | hsa_circ_0005232 | SLC8A1 | **circSlc8a1** | chr17 | 82047148 | 82048978 | mmu_circ_0000823 | Slc8a1 |
| **circTULP4** | chr6 | 158733082 | 158735300 | hsa_circ_0131202 | TULP4 | **circTulp4** | chr17 | 6137210 | 6139156 | mmu_circ_0000723 | Tulp4 |
| **circUNC79** | chr14 | 93934016 | 93963632 | hsa_circ_0102954 | UNC79 | **circUnc79** | chr12 | 104229559 | 104260086 | mmu_circ_0000411 | Unc79 |
| **circZNF609** | chr15 | 64791491 | 64792365 | hsa_circ_0000615 | ZNF609 | **circZfp609** | chr9 | 65642428 | 65643302 | mmu_circ_0001797 | Zfp609 |
| **circZNF827** | chr4 | 146744573 | 146770713 | hsa_circ_0001447 | ZNF827 | **circZfp827** | chr8 | 81642073 | 81660562 | mmu_circ_0001698 | Zfp827 |

Table S3

| dishRNA-circZNF827 vs. dishRNA-Irr<br>+RA<br>fold change>2, p<0.05 | | dishRNA-circZNF827 vs. dishRNA-Irr<br>-RA<br>fold change>2, p<0.05 | | dishRNA-circZNF827 vs. dishRNA-Irr<br>+RA, BrU IP<br>fold change>2, p<0.05 | |
|---|---|---|---|---|---|
| *Upregulated genes* | *Downregulated genes* | *Upregulated genes* | *Downregulated genes* | *Upregulated genes* | *Downregulated genes* |
| NGFR | NQO1 | FN1 | NTNG1 | NQO1 | ATP8A2 |
| SYT13 | NTS | NQO1 | FOS | NTS | NGFR |
| NTRK1 | FN1 | TNR | GRM8 | FN1 | MYC |
| | BNIP3 | TNC | SNCAIP | TNR | SYT13 |
| | TNR | DDC | CDKN1A | TNC | PMP22 |
| | NEGR1 | BNIP3 | FAS | TAF4B | |
| | TBPL1 | NEGR1 | LRP1 | NELL2 | |
| | TNC | STAT3 | PLA2G16 | FXN | |
| | PTEN | | ACHE | NEGR1 | |
| | STAT3 | | DCX | | |
| | GTF2A1 | | CADPS | | |
| | JAM3 | | ARC | | |
| | OXR1 | | PRKCE | | |
| | CHRNA7 | | STX1B | | |
| | PPT1 | | | | |
| | MMP16 | | | | |
| | MYH10 | | | | |
| | PLCB4 | | | | |
| | NELL2 | | | | |
| | GALC | | | | |
| | CHMP2B | | | | |
| | PLS1 | | | | |
| | SORL1 | | | | |
| | LPAR1 | | | | |
| | GSTP1 | | | | |
| | MGMT | | | | |
| | PGK1 | | | | |
| | EIF2S1 | | | | |
| | IDE | | | | |

### Table S4

| | | |
|---|---|---|
| **Oligonucleotides for construction of pFRT/U6-dishRNA** | | |
| | Sense (5’-3’) | Antisense (5’-3’) |
| hscircTULP4 | GATCACAAATCTCGCTATTGTGGCCCCACAATAGCGAGATTTGCTTTTT | TCGAAAAAGCAAATCTCGCTATTGTGGGGCCACAATAGCGAGATTTGT |
| hscircZNF609 | GATCACAACATCATTGCTTTTCAGACGAAAAGCAATGATGTTGCTTTTT | TCGAAAAAGCAACATCATTGCTTTTCGTCTGAAAAGCAATGATGTTGT |
| hscircSLC8A1 | GATCAGTCACAACCTAACAATTTCATAATTGTTAGGTTGTGACCTTTTT | TCGAAAAAGGTCACAACCTAACAATTATGAAATTGTTAGGTTGTGACT |
| hscircHDGFRP3 | GATCAAGTTCATCAATCCCTTCACTGGAAGGGATTGATGAACTCTTTTT | TCGAAAAAGAGTTCATCAATCCCTTCCAGTGAAGGGATTGATGAACTT |
| hscircCDYL | GATCAATCCTTTCAACCTTTCCCGTTGGAAAGGTTGAAAGGATCTTTTT | TCGAAAAAGATCCTTTCAACCTTTCCAACGGGAAAGGTTGAAAGGATT |
| hscircCAMSAP1 | GATCAAGGGATGTTATCTTGTTGATCAACAAGATAACATCCCTCTTTTT | TCGAAAAAGAGGGATGTTATCTTGTTGATCAACAAGATAACATCCCTT |
| hscircZNF827 | GATCAGTAGCTTTCCGCTGGAGCTGACTCCAGCGGAAAGCTACCTTTTT | TCGAAAAAGGTAGCTTTCCGCTGGAGTCAGCTCCAGCGGAAAGCTACT |
| hscircANKIB1 | GATCATCTTTCACATTCATGAGCTCGCTCATGAATGTGAAAGACTTTTT | TCGAAAAAGTCTTTCACATTCATGAGCGAGCTCATGAATGTGAAAGAT |
| hscircUNC79 | GATCACTTGGAAGCAACTGTGTACTGACACAGTTGCTTCCAAGCTTTTT | TCGAAAAAGCTTGGAAGCAACTGTGTCAGTACACAGTTGCTTCCAAGT |
| mmcirczfp827 | GATCAGTAGCTTTCCGCTGGAGCAGACTCCAGCGGAAAGCTACCTTTTT | TCGAAAAAGGTAGCTTTCCGCTGGAGTCTGCTCCAGCGGAAAGCTACT |
| hnRNPK #1 | GATCAATAATTCCTCCTGCTAGACTCCTAGCAGGAGGAATTATCTTTTT | TCGAAAAAGATAATTCCTCCTGCTAGGAGTCTAGCAGGAGGAATTATT |
| hnRNPK #2 | GATCATTCATAAGCCATCTGCCATTCGGCAGATGGCTTATGAACTTTTT | TCGAAAAAGTTCATAAGCCATCTGCCGAATGGCAGATGGCTTATGAAT |
| hnRNPL #1 | GATCAATTTCACCTTCTCCACATTGCTGTGGAGAAGGTGAAATCTTTTT | TCGAAAAAGATTTCACCTTCTCCACAGCAATGTGGAGAAGGTGAAATT |
| hnRNPL #2 | GATCACACTTTTGCCTGAGAATACTTATTCTCAGGCAAAAGTGCTTTTT | TCGAAAAAGCACTTTTGCCTGAGAATAAGTATTCTCAGGCAAAAGTGT |
| Irr | GATCATCGTCATAACGTTCATAGGCGTATGAACGTTATGACGACTTTTT | TCGAAAAAGTCGTCATAACGTTCATACGCCTATGAACGTTATGACGAT |
| **Primers for construction of pCCL/U6-dishRNA-PGK-eGFP-MCS** | | |
| | Sense (5’-3’) | Antisense (5’-3’) |
| U6-dishRNA | AAAATCGATAAGGTCGGGCAGGAAGAGGG | AAACGTACGAAACGGGCCCTCTAGACTCG |
| **Primers for construction of pcDNA3/circRNA** | | |
| | Sense (5’-3’) | Antisense (5’-3’) |
| circZNF827 | AAAGGATCCCGGAAAGCTACAAGGAAACAC | AAAGCGGCCGCCTGGAGCTGATATTCTTCCG |
| circTULP4 | AAAGGATCCGATTTGTAAGACTCCAGGGCCTCCC | AAAGCGGCCGCCTCGCTATTGTGGCCCCGGAGGTTG |
| circZNF609 | AAAAAGCTTGCAATGATGTTGTCCACTGG | AAAGCGGCCGCCTTTTCAGACTTGACTTTCTTTAG |
| circHDGFRP3 | AAAGGATCCATTGATGAACTCCCAGAGGG | AAAGCGGCCGCCCCTTCACTGGTTTTCTGCAAG |
| circSLC8A1 | AAAGGATCCGTTGTGACAGTTGGAAGTGTC | AAAGCGGCCGCACAATTTCATCATTCTGGA |
| **Primers for construction of pcDNA3.1(+)-Laccase2-circRNA** | | |
| | Sense (5’-3’) | Antisense (5’-3’) |
| circZNF827 | AAATTAATTAACGGAAAGCTACAAGGAAACAC | AAACCGCGGCTGGAGCTGATATTCTTCCG |
| **Primers for construction of pcDNA5_FRT/TO-FLAG-RBP** | | |
| | Sense (5’-3’) | Antisense (5’-3’) |
| hnRNPU | ATTAGGATCCATGAGTTCCTCGCCTGTTAA | ATTAGCGGCCGCTCAATAATATCCTTGGTGAT |
| DHX9 | ATTAGGATCCATGGGTGACGTTAAAAATTTTC | ATTAGCGGCCGCTTAATAGCCGCCACCTCCTC |
| DDX3X | ATTAGGATCCATGAGTCATGTGGCAGTGGA | ATTAGCGGCCGCTCAGTTACCCCACCAGTCAA |
| hnRNPK | AAAGGTACCATGGAAACTGAACAGCCAGA | AAAGCGGCCGCTTAGAATCCTTCAACATCTG |
| hnRNPL (short isoform) | AAAGGATCCATGTCGCGGAGGCTGCTGCC | AAAGCGGCCGCTTAGGAGGCGTGCTGAGCAG |
| hnRNPL (long isoform) | AAAGGATCCATGCCTAAAAAGAGACAAGC | AAAGCGGCCGCTTAGGAGGCGTGCTGAGCAG |
| **Primers for RT-qPCR** | | |
| | Sense (5’-3’) | Antisense (5’-3’) |
| mmcircMed13L | CCAACCTGCTGTGTGTATGG | TGCCCTCCAAAATTGTACCT |
| mmcircMagi1 | TAGTCCACCCGGAGAATGAG | CTTCATAGGTGCCGACTTCC |
| mmcircHdgfrp3 | TGACACGAGAAACACGACTG | ACTTGTTTGCTGGAGGCTTC |
| mmcircEzh2 | TACAGCCTGTGCACATCATG | TTTTACACGCTTCCGCCAAC |
| mmcircAnkib1 | TATGCTGCCTTAGACAAGCG | AGTGGCACGTTCTTTCTTCC |
| mmcircAff3 | CAAGCTCCTGACGAATCTCC | CTGATGCCTGAACTTCCAGA |
| mmcircNfix | TGTCCAGCCACATCACATTG | TTTGACATCCGCTTCTCGTG |
| mmcircRmst | GAAGGATCTGCCAAGCTCAC | CACTTCGTCCCTGACTCCAT |
| mmcircSlc8a1 | TCTGGAGCTCGAGGAAATGT | TTGGGTGGGAGACTTAATCG |
| mmcircTulp4 | AGAAACCTGTGTGCAGAAGG | TTCATGACTGGAAGGTCCTG |
| mmcircZfp609 | CCGGCCACTAAAGAAAGTCA | AGTCAACGTCCCACCTCAAG |
| mmcircZfp827 | AAAAGATCTGCTGCCTGGTG | AGTGCTGTAGCCAATGTTCC |
| mmNanog | CCTCCAGCAGATGCAAGAACTC | CTTCAACCACTGGTTTTTCTGCC |
| mmNestin | GAAGGTGGGCAGCAACTGGCA | TCAGCCTCCAGCAGAGTCCTGT |
| mmTrkB | AGCAGCCCTGGTATCAGCTA | CTTGATGTTCTTCCGGGTGT |
| mmB-Tub III | ACTTGGAACCTGGAACCATGG | GGCCTGAATAGGTGTCCAAAGG |
| mmGAPDH | TGGGAAGCTTGTCATCAACG | TTTTGGCTCCACCCTTCAAG |
| hscircTULP4 | CTGTGTGCAGGAGACGCTAC | CTTCCCCAGTCATTTCAGGA |
| hscircZNF609 | AGGAAGGGGAGAATGAGTGT | GTTCTCAGACCTGCCACATT |
| hscircSLC8A1 | GAGGGGAGGATTTTGAGGAC | AGACTTAATCGCCGCATGTT |
| hscircHDGFRP3 | GCAACGACACAAGAAACACA | GTTTCATGGGTGCCAAAA |
| hscircCDYL | GGCCCAGGATACACCCACTA | AGCCTTTCCACCGAACCAAA |
| hscircCAMSAP1 | GATGATCAGCATCGAGAAGG | ATAACAGGCGGCTTAATGTG |
| hscircZNF827 | TCACCATTCACCTCCAATTC | AGTGCTGTAGCCAACATTCC |
| hscircANKIB1 | AGGCCTCAGGATCTTCGTAG | CACTTCTGGAACACTTCTGCA |
| hscircUNC79 | CGTATGGGCCTTCACAAGTG | TCCTGCAAGTACCGGATCTT |
| hsRAR$\alpha$ | CTGCCTGGACATCCTGATCC | CAAAGACCAGGTCGGTGAGG |
| hsRAR$\beta$ | GAAGGAGACTTCGAAGCAAG | ATGGTCAGCACTGGAATTCG |
| hsRAR$\gamma$ | TCGCTAGAGGCATTGGGGTG | GGGCATCAGCACTAAGGGAG |
| hsTUBB3 | GGCCTCTTCTCACAAGTACG | CCACTCTGACCAAAGATGAAA |
| hsTrkB | ACCACTGGTGCATTCCATTC | CCCCATTGTTCATGTGAGTG |
| hsNEFL | GAGTGAAATGGCACGATACC | TAGCCACTGGTTATGCTTCC |
| hsMAP2 | CAAAGAGAATGGGATCAACG | CAGGACTGCTACAGCCTCAG |
| hsGAPDH | GTCAGCCGCATCTTCTTTTG | GCGCCCAATACGACCAAATC |
| hs$\beta$-actin pre-mRNA | TTCCCCAGTGTGACATGGTG | ACCATGTCACACTGGGGAAG |
| hsNGFR | CCGAGGCACCACCGACAACC | GGGCGTCTGGTTCACTGGCC |
| hsc-FOS | TCGCCTGTCAACGCGCAGG | GACTGGTCGAGATGGCAGT |
| hsZNF827 | ACCCGGATGTTTGAGTGTGA | ACCACTGTCCTGAGTTGAGC |
| hshnRNPL | ACAAACCCCAATCTCAGTGG | CATATTCTGCGGGGTGATCT |
| hsNGFR promoter | CTGATAGGGGGCAGTTAGGG | GCCACAGGGTTCACTCTCTT |
| hsGAPDH promoter | Pierce Magnetic ChIP Kit (Thermo Scientific) | Pierce Magnetic ChIP Kit (Thermo Scientific) |
| **Probes for Northern blot (5’-3’)** | | |
| circZNF827 | TCACCATCTGTGTTTCCTTGTAGCTTTCCGCTGGAGCTGATATTCTTCCGTGCAGCACGG | |
| $\beta$-actin | AATGTCACGCACGATTTCCCGCTCGGCCGTGGTGGTGAAGCTGTAGCCGC | |

### Table S5

| Antibody | Manufacturer | Catalog number |
|---|---|---|
| BrU | BD Biosciences | 555627 |
| DDX3X | Sigma-Aldrich | HPA00648 |
| FLAG | Sigma-Aldrich | F7425 |
| GAPDH | Invitrogen | AM4300 |
| hnRNP C1/C2 | Kind gift form Seraphin Pinol-Roma | |
| hnRNPK | Abcam | ab39975 |
| hnRNPK | Santa Cruz Biotechnology | sc-28380 |
| hnRNPL | Abcam | ab6106 |
| hnRNPU | Sigma-Aldrich | HPA058707 |
| HuR | Santa Cruz Biotechnology | sc-5261 |
| IgG | Cell Signaling | 2729S |
| LARP1 | Cell Signaling | 70180S |
| MAP2 | Santa Cruz Biotechnology | sc-390543 |
| NGFR | Cell Signaling | 8238S |
| RNA Pol II | Thermo Scientific (Pierce Magnetic ChIP Kit) | 26157 |
| TUBB3 | Cell Signaling | 8572S |
| ZNF827 | Santa Cruz Biotechnology | sc-514943 |
